# Unraveling interindividual differences and functional consequences of gut microbial metabolism of immunosuppressants

**DOI:** 10.1101/2024.03.28.586928

**Authors:** Maral Baghai Arassi, Nicolai Karcher, Eleonora Mastrorilli, Matthias Gross, Amber Brauer-Nikonow, Raymund Hackett, David Czock, Burkhard Tönshoff, Georg Zeller, Michael Zimmermann

**Affiliations:** Department of Pediatrics I, University Children’s Hospital Heidelberg; Heidelberg, Germany; Structural and Computational Biology Unit, European Molecular Biology Laboratory; Heidelberg, Germany; Leiden University Center for Infectious Diseases (LUCID), Leiden University Medical Center; Leiden, Netherlands; Department of Clinical Pharmacology and Pharmacoepidemiology, Heidelberg University Hospital; Heidelberg, Germany

## Abstract

A major challenge in kidney transplantation (KT) is the large interpatient variability in the pharmacokinetics of immunosuppressive drugs. Here, we explored the role of the gut microbiome in interindividual variation in immunosuppressive drug metabolism. Analysis of 38 fecal communities, including 10 from KT recipients, and 45 bacterial species against 25 drugs, revealed significant interindividual and drug-specific differences in metabolism. Notably, 15 of 16 immunosuppressants tested were metabolized by at least one microbial community, and we found specific bacterial species, such as *Bacteroides uniformis*, to be potent metabolizers. We identified 18 different metabolites for 16 drugs, including two previously undescribed metabolites for sirolimus and everolimus. Our study reveals the functional impact of microbial metabolism on key immunosuppressants, including inactivation of tacrolimus, activation and potential increase in toxicity of mycophenolate mofetil (MMF), and shows that the microbial metabolite of methylprednisolone exhibits a 2.6-fold increase in epithelial permeability compared to the parent drug. Through a gain-of-function genetic screen we identified the *B. uniformis* enzyme BACUNI_RS05305 to be responsible for MMF activation. Using machine learning to model microbial community drug metabolism, abundance features of prevalent species predicted the biotransformation of some drugs well, while for others, a priori experimental information on bacterial genes and enzyme protein structures led to improved predictions. Our research highlights the potential of gut microbiome features to explain interindividual variability in immunosuppressive therapies and sets the stage for clinical trials to identify microbiome-encoded signatures predictive of drug metabolism in KT patients.

## INTRODUCTION

Kidney transplantation is an important therapy for end-stage organ disease (*1, 2*). The introduction of potent immunosuppressants, such as the calcineurin inhibitor tacrolimus and the proliferation inhibitor mycophenolate mofetil (MMF), has improved the clinical outcome of kidney transplantation (KT) making them a cornerstone of modern transplantation medicine. However, the large interpatient variability in pharmacokinetics and pharmacodynamics represent a major challenge in clinical practice (*3, 4*). Underexposure of immunosuppressants may lead to acute graft rejection, whereas overexposure bears the risk of serious drug-related side effects, such as infections, nephrotoxicity, neurotoxicity and metabolic disorders (*5*). Despite decades of research aiming to predict an individual’s drug pharmacokinetics, known biomarkers, such as cytochrome P450 polymorphisms, have limited accuracy (*6*). Therefore, novel insights into the factors that determine the pharmacokinetics and pharmacodynamics of immunosuppressants are urgently needed to optimize personalized immunosuppression in SOT.

One factor in drug metabolism that has recently garnered attention is the human gut microbiome (*7–10*). Bacteria, the major constituent of the complex gastrointestinal ecosystem, harbor a myriad of metabolic enzymes with a tremendous potential to metabolize drugs. We have previously found that a substantial fraction of drugs, including some widely used immunosuppressants are metabolized by human gut bacteria (*7*). The remarkable interindividual genetic variability of human gut microbiomes, far surpassing that of human genomes, suggests that the microbiome may partially contribute to differences in drug exposure and response (*11*). This is particularly relevant in the context of drugs with large interpersonal variability, such as immunosuppressants (*5*). Moreover, research indicates that gut microbial modifications of immunosuppressants can influence drug elimination rates, toxicity (*12*) and efficacy (*13*). Despite these insights, a comprehensive understanding of the interindividual differences, functional consequences and underlying mechanisms of gut microbial metabolism of immunosuppressants is still lacking.

To close this knowledge gap, we assayed 25 medical drugs commonly used in solid organ transplantation, with a special focus on KT, including 16 immunosuppressants for their metabolism by 38 distinct human-derived gut microbial communities and 45 representative gut bacterial species. These experiments revealed pronounced interindividual and drug-specific differences in microbiota drug metabolism. We further observed that microbial metabolism of the three key immunosuppressants used in KT – tacrolimus, MMF, and methylprednisolone, can cause drug inactivation, prodrug activation, increased toxicity, and altered intestinal absorption. Based on these data and employing genome-wide functional screens, we identified mechanistic determinants of microbial drug metabolism, including specific bacterial species and drug-metabolizing bacterial enzymes. These findings provide the basis for predicting interindividual differences in drug metabolism from microbiome features.

## RESULTS

### Subhead 1: Pronounced interindividual differences in microbial metabolism of immunosuppressants

To systematically assess potential interindividual differences in gut microbial metabolism of immunosuppressive drugs, we analyzed the drug-metabolizing capacity of human gut microbial communities derived from 19 individual donors, including 14 healthy adults and children and 5 adult kidney transplant recipients (Fig. 1A, Table S1). The stool donors were selected to provide a comprehensive representation of both the general population and transplant recipients in particular. Characterizing the microbial composition through metagenomic sequencing, we found them to be compositionally diverse and representative compared to a large collection of human gut microbiomes (n = 3189, PERMANOVA R2 0.138%, see Fig. 1B, S1C and Table S2 for details). Drug incubation under anaerobic and microaerobic (5% O_2_) conditions, to mimic differences in oxygen availability along the gastrointestinal tract (*14*), yielded 38 tested human-derived gut microbial communities (Fig. 1B, see S1A, B and Table S3, S4 for details). We assayed the metabolism of each of the microbial communities against each of the 25 drugs, including 16 common immunosuppressants, as well as the cornerstone immunosuppressants tacrolimus, MMF and methylprednisolone, which are known for their interindividual pharmacokinetic variability. Additionally, we incorporated 9 non-immunosuppressive drugs frequently used as comedication following kidney transplantation (Fig. 1C, Table S5). In order to test the metabolic potential of microbial communities *ex vivo* we incubated each of the 475 community-drug combinations in quadruplicate and measured drug metabolism kinetics (8 timepoints) over 12 hours by liquid chromatography-coupled mass spectrometry (LC–MS, Fig. 1A). Of the 16 immunosuppressants tested, 15 were significantly eliminated after incubation (>25 % reduction in abundance, FDR-corrected P value ≤0.05) by at least one of the tested gut microbial communities compared to no-bacteria controls. Gut microbial communities differed widely in their ability to metabolize drugs with respect to both the number of immunosuppressants metabolized and the extent of metabolism of individual drugs (Fig. 1D, Table S6 and S7). MMF was metabolized by the highest number of gut microbial communities (32/38, 84%) followed by glucocorticoids, such as prednisolone (29/38, 76%) and methylprednisolone (26/38, 68%) under both anaerobic and microaerobic conditions. Additionally, we observed significant microbial metabolism of tacrolimus by 21% of the tested gut microbial communities (8/38). Altogether, we found substantial microbial metabolism of the most clinically relevant immunosuppressants with pronounced interindividual differences.

**Fig. 1.**
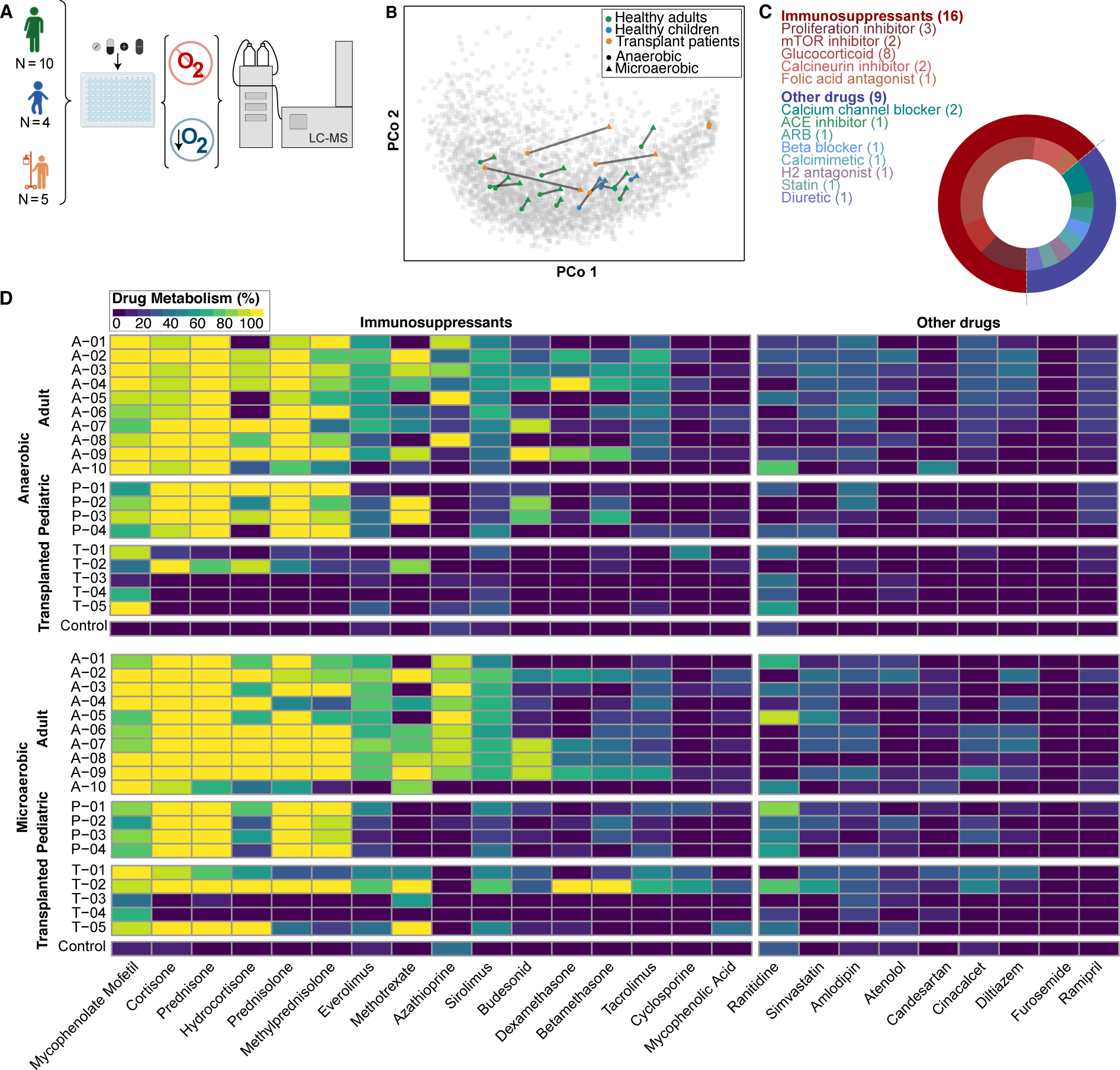
Drug metabolizing activity of human-derived gut microbial communities. **(A)** Schematic of the assay. **(B)** Principal Coordinates Analysis (PCoA) comparing the composition of 38 human-derived gut microbial communities, cultivated under anaerobic conditions for 8 hours before sequencing, to 3189 reference human gut microbial metagenomes from 10 datasets in industrialized countries (corresponding to the Westernized control samples used in (*30*), Table S2). Circles and triangles denote the tested communities, color-coded by population source (see inset). Gray circles represent samples from reference metagenomes. Colored circles indicate samples cultivated anaerobically, triangles represent microaerobically cultivated samples. Gray lines connect *ex vivo* cultures originating from the same stool sample. **(C)** Overview of tested drug classes, with the number of representatives tested indicated in brackets. Shades of red represent immunosuppressants, while shades of blue denote other, non-immunosuppressive drugs. Pie chart visualizes the proportion of each drug class among all drugs tested. **(D)** Heat map depicting the metabolism of the 25 tested drugs by 38 human-derived gut microbial communities. The communities are arranged according to the population source (adults, children, transplant patients). Drugs are ordered based on drug class (immunosuppressant vs. other drugs) and observed metabolic activity.

### Subhead 2: Identification of immunosuppressant-metabolizing gut bacterial species

Given the large interindividual differences of gut microbial metabolism of immunosuppressants, we next aimed to identify specific bacterial isolates capable of performing immunosuppressant biotransformation. To this end, we assessed the metabolic activity of 45 selected gut bacterial species (Table S8) against our panel of 25 drugs. The selected species encompass the five major phyla found in the human gut and represent on average 43.9% of the total microbial abundance of the 38 human-derived microbial communities tested (see Fig. S2 for details). Similarly, they were highly abundant in our reference human gut microbial metagenomes with a mean relative abundance of up to 10% (Fig. 2). We additionally assayed those 18 of the 45 (40%) bacterial species under microaerobic conditions, that are commonly found in the upper gastrointestinal tract, including the small intestine. We found that single-species bacterial cultures metabolized immunosuppressants under both anaerobic and microaerobic conditions (Fig. 2, Table S9, 10). Metabolized drugs included positive controls expected to be metabolized by gut bacteria, such as diltiazem by *Bacteroides thetaiotaomicron* (*7*) or glucocorticoids by *Clostridium scindens* (*7, 15*). The overall number of metabolized drugs was higher under anaerobic compared to microaerobic conditions (20 vs 8), but this may be due to the higher number of species tested under anaerobic compared to microaerobic conditions (44 versus 18 species). MMF, azathioprine and sirolimus were most extensively metabolized under both culture conditions. Overall, these results indicate strong microbial metabolism of clinically relevant immunosuppressants with pronounced species-specific differences. Furthermore, these results align well with our observed drug metabolism by gut microbial communities, as 16 out of the 25 tested drugs were metabolized by both gut microbial communities and species. This hence provides the basis to understand drug metabolism of gut microbial communities as a functional trait of its members.

**Fig. 2.**
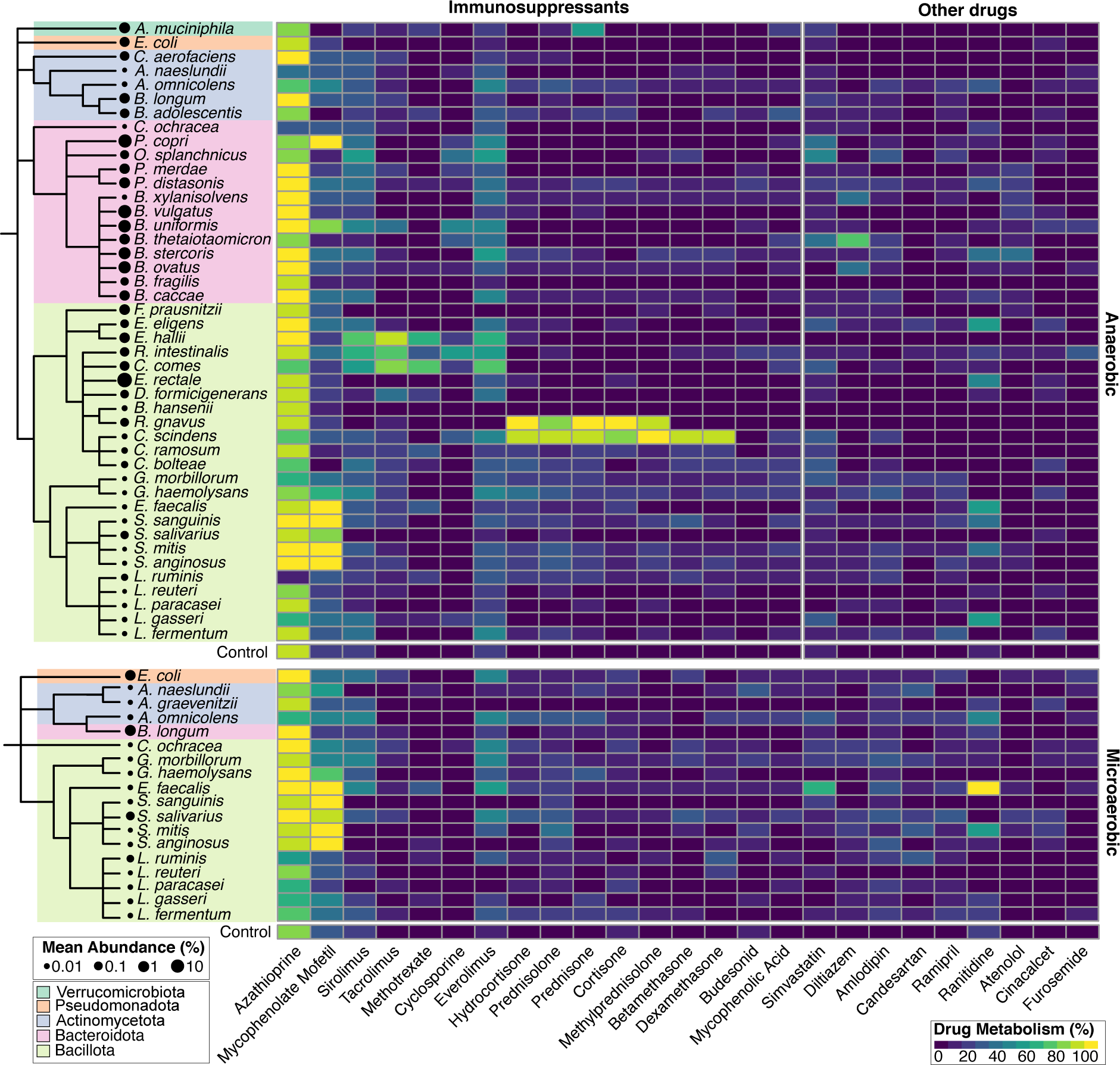
Drug metabolizing activity of individual gut bacterial species. Heat map illustrating the metabolism of the 25 tested drugs by 45 individual gut bacterial species. The species are arranged taxonomically, with dendrogram color coding representing the respective phylum. Black dots indicate the mean abundance of each species estimated on the reference metagenomes (see Fig. 1B). Drugs are ordered based on drug class (immunosuppressant vs. other drugs) and observed metabolism.

### Subhead 3: Identification of microbial drug metabolites

To identify bacteria-produced drug metabolites we leveraged insights from our prior high-throughput screen (*7*) and existing literature (*16, 17*). Mining untargeted metabolomics data, we identified a total of 18 different drug metabolites for 16 different drugs (Table S11 and Fig. 3A). The accumulation of a given drug metabolite varied between individuals and species (Table S12, 13). Intriguingly, the accumulation of drug metabolites only partially correlated with the depletion of respective parent compounds, suggesting that additional drug metabolites may be produced (Fig. 3A, S3). Building on the established keto-reduction pathway that deactivates tacrolimus (*13*) and structural similarities with sirolimus, and everolimus, we identified 9-hydroxy-everolimus (982.587 Da) and 9-hydroxy-sirolimus (938.5606 Da) as two new drug metabolites (Fig. 3B-D). This finding was corroborated by the fact that the same three microbial communities were consistently the most effective in generating the respective microbial metabolite of tacrolimus, sirolimus and everolimus (Fig. 3E). Moreover, the detection of both metabolites was confirmed in our single-species screen (Fig. S3 and Table S14, S15 for details). Taken together, the identification of 18 metabolites, including two previously unknown ones, coupled with the observed interpersonal variability in metabolite production, not only enhance our understanding of microbial drug interactions but also opens avenues for further research into the functional implications of gut bacterial drug metabolism.

**Fig. 3.**
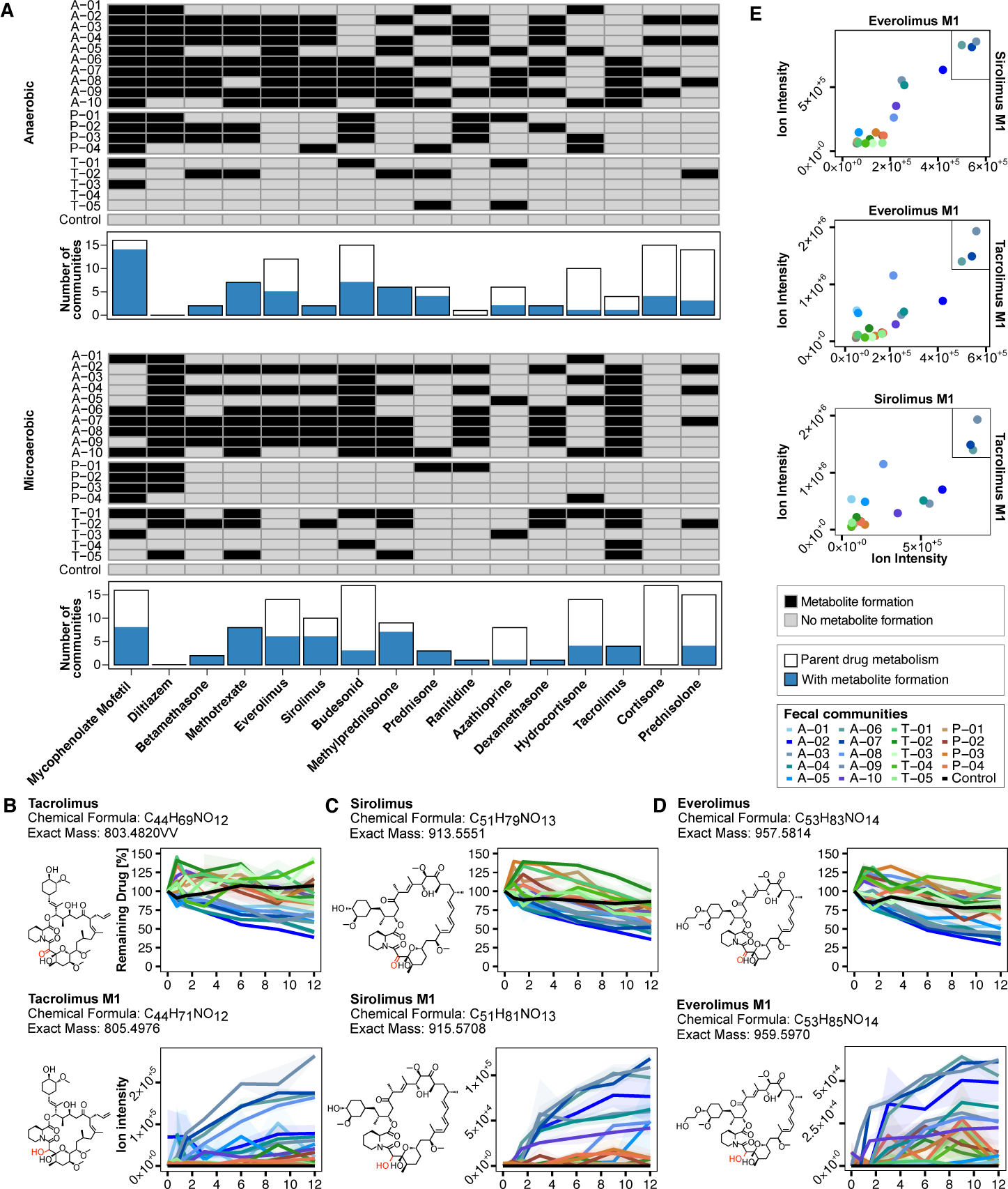
Identified drug metabolites in human-derived gut microbial communities. **(A)** Heat map illustrating microbial drug metabolite formation based on the parent drug (columns). Bar outlines below the heat map depict the number of communities metabolizing a given parent drug. Blue filled bars represent the number of communities where additional metabolite formation was observed. Drugs are sorted based on metabolite formation, while microbial communities are sorted based on the source population. **(B-D)** Chemical structure and exact mass of parent drugs and their respective metabolites for tacrolimus, sirolimus, and everolimus. Line plots depict the degradation of the parent drug and formation of metabolites over time under anaerobic conditions, with individual communities color-coded. Lines and shaded areas indicate the mean and s.e. of n=4 assay replicates, respectively. **(E)** Scatterplots depict pairwise correlation of metabolite formation of tacrolimus, everolimus, and sirolimus by human gut microbial communities under anaerobic conditions, with color-coded circles representing individual communities. In the upper right panel of each plot a black box highlights the same top three communities most actively forming the metabolite.

### Subhead 4: Intestinal transport of microbiota-produced metabolites

To gain insights into the influence of microbial drug metabolism on enteric absorption, a key determinant of systemic drug distribution, we used an *in vitro* approach to assess transport rates of drugs and drug metabolites across a gut epithelial monolayer (*18*). We focused on methylprednisolone and its bacteria-produced metabolite hydroxymethylandrostadienedione (HMADD, Fig. 4A). This selection was motivated by the pivotal clinical role of methylprednisolone within the first-line immunosuppressive regimen in KT and its notable variability in gut microbial metabolism (Fig. 4A). To establish an epithelial monolayer, Caco-2 cells were seeded into transwells and methylprednisolone as well as its bacteria-produced metabolite were added to the apical epithelial side after the establishment of a stable Caco-2 cell monolayer (Fig. 4C). This setup enabled independent access to both sides of the monolayer with each compound incubated in quadruplicate and its transport kinetics measured every 30 minutes over a time period of 5 hours using LC–MS. This analysis revealed that bacteria-produced HMADD exhibited different absorption characteristics compared to its parent compound (Fig. 4D, Table S16). To control for the potential impact of bacterial culture supernatant on the epithelial barrier, we also assayed the epithelial absorption of the pure chemical standard of both the parent drug and its metabolite which confirmed our observation of faster epithelial absorption of HMADD compared to methylprednisolone (Fig. S4). Indeed, quantification of the effective permeability (P_eff_) (*18*) revealed that the microbial methylprednisolone metabolite has a 2.6-fold higher permeability (8.11 cm/sec) compared to the parent drug (3.11 cm/sec, p=0.029, see also Fig. S4 for details). Collectively, these findings suggest that microbiota-produced metabolites, such as HMADD, may exhibit distinct absorption rates compared to their parent drugs across gut epithelial barriers, implying the possibility of systemic absorption from the large intestine, potentially impacting drug pharmacokinetics.

**Fig. 4.**
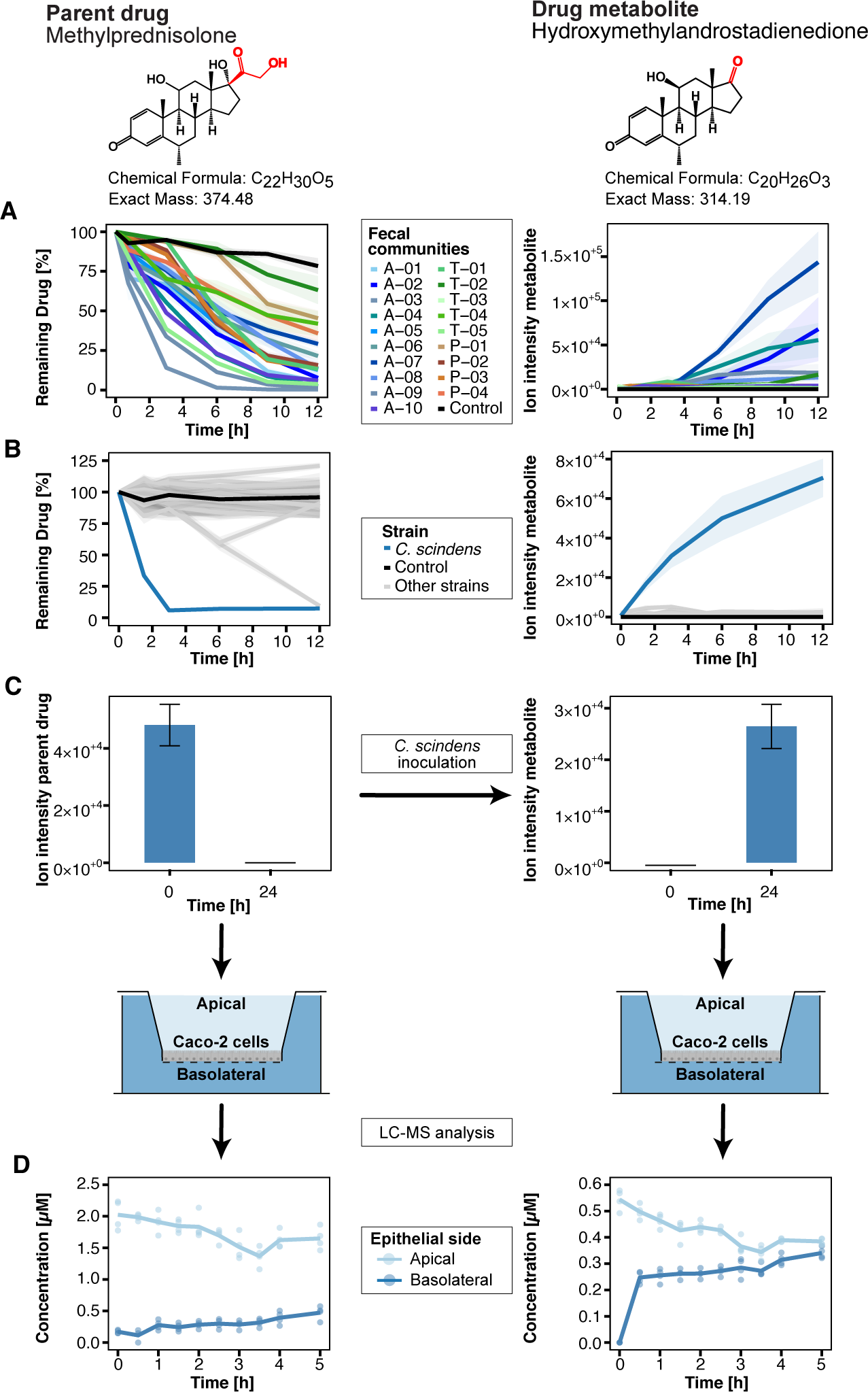
Epithelial transport characteristics of methylprednisolone and its microbial metabolite. **(A)** Gut microbial community metabolism of methylprednisolone (left) and formation of its microbial metabolite hydroxymethylandrostadienedione (right) under anaerobic conditions over time. Individual gut microbial communities are represented by color-coded lines. Lines and shaded areas depict the mean and s.e. of n=4 assay replicates, respectively. **(B)** Metabolism of methylprednisolone and formation of hydroxymethylandrostadienedione over time by individual bacterial species under anaerobic conditions. *C. scindens* used for metabolite accumulation shown in (C) is depicted by a blue line, while the other 44 species are shown with gray lines. Lines and shaded areas depict the mean and s.e. of n=4 assay replicates, respectively. **(C)** Barplot showing the ion intensity of methylprednisolone at the start of the *C. scindens* inoculation vs. after 24 hours, with metabolite buildup on the right. Barplots and error bars represent the mean and standard error of n=6 assay replicates. The lower panel includes a schematic of the transport assay, with light blue depicting the apical side of the epithelial Caco-2 monolayer and dark blue the basolateral side. **(D)** Line plot showing the concentration over time of methylprednisolone and its microbial metabolite hydroxymethylandrostadienedione on the apical (light blue) and basolateral side (dark blue), respectively. Lines represent the mean of n=4 assay replicates while circles represent individual measurements.

### Subhead 5: Identification of immunosuppressant-metabolizing bacterial enzymes

To identify the genetic determinants of gut bacterial drug metabolism, we focused on MMF due to reported gastrointestinal toxicity attributed to its active moiety, MPA (*12*). Intriguingly, we found that MMF is metabolized by 16 different microbial communities (Fig. 5A) and 23 individual bacterial species under anaerobic conditions including the strong MMF-metabolizer *Bacteroides uniformis* (Fig. 5B). To elucidate the genetic basis of gut microbial drug metabolism we employed a previously established gain-of-function approach using *B. uniformis* (*7*). In brief, we extracted and sheared genomic DNA (gDNA) into 2-8 kb fragments. gDNA fragments were then cloned into an *Escherichia coli* expression vector. Transformation and arraying of individual clones yielded 26,000 transformed *E. coli* clones with an average insert size of 3 kb (Fig. 5C), which translates into a 15-fold genome coverage of *B. uniformis*. Through consecutive pooling of clones (*7*) and mass spectrometry-based testing for MMF metabolism, we identified BACUNI_RS05305 (Fig. 5D), a microbial esterase, as an MMF-metabolizing enzyme. To validate these findings, we expressed the enzyme encoded by BACUNI_RS05305 in *E. coli* and demonstrated its MMF-metabolizing capability (Table S17). Furthermore, we showed that deleting BACUNI_RS05305 in *B. uniformis* abolished MMF activation and that complementation with promoters of different strength restored MMF metabolism of the BACUNI_RS05305 knock-out strain (Fig. 5E). Additionally, we investigated BACUNI_RS05310, a neighboring gene which exhibits 85% sequence similarity to RS05305. Deleting BACUNI_RS05310 did not affect MMF metabolism (Fig. S5), indicating that subtle structural variations in enzymes, undetectable by sequence analysis alone, could influence enzyme functionality. In summary, our study has successfully identified the gene BACUNI_RS05305 as a key player in the activation of the immunosuppressive prodrug MMF to its active form MPA, shedding light on the genetics of this gut microbial drug metabolization and suggesting molecular markers that could be used to explain interindividual variability in drug metabolism.

**Fig. 5.**
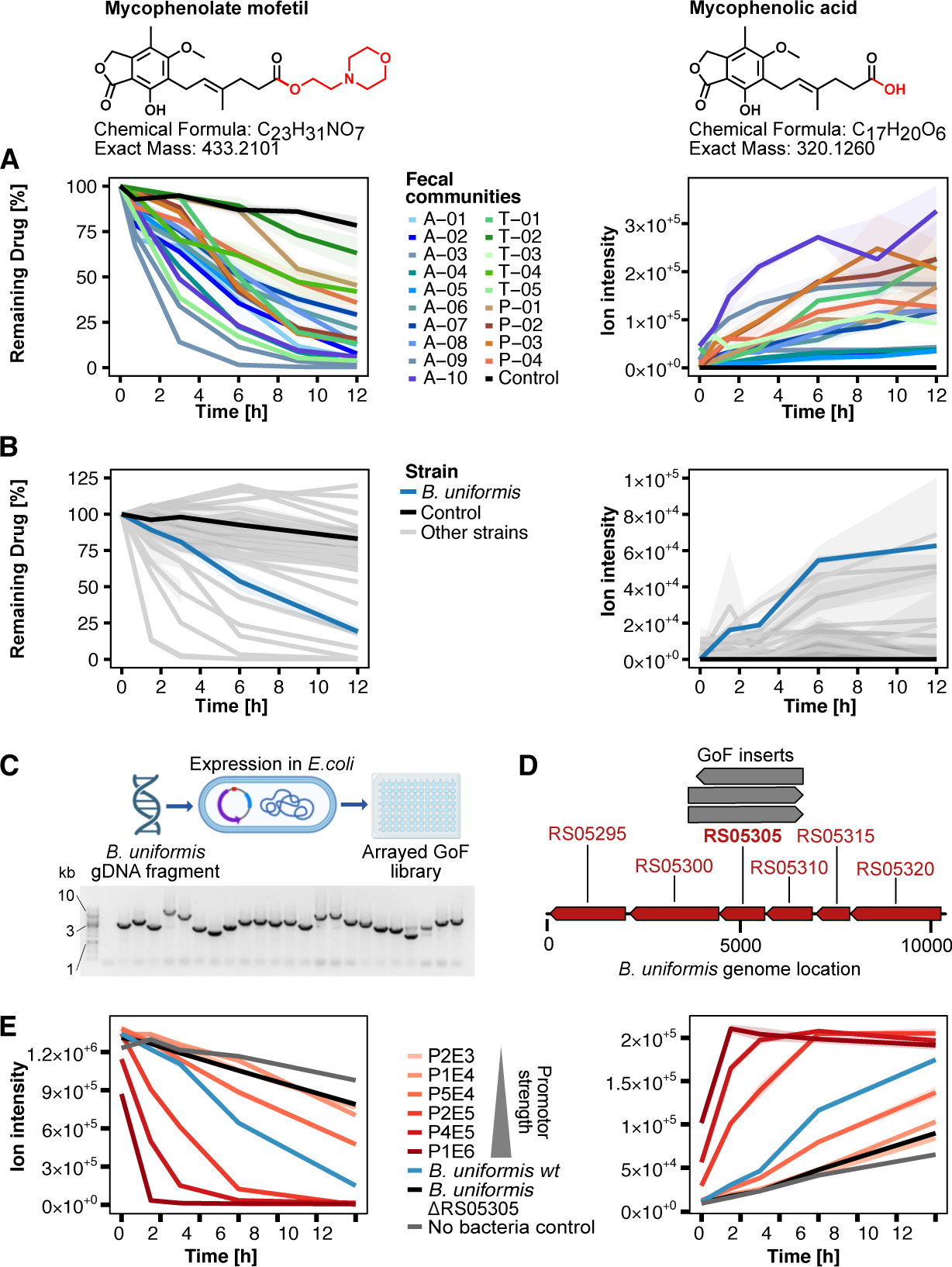
Identification of MMF-metabolizing bacterial enzyme. **(A)** Gut microbial community metabolism of mycophenolate mofetil (MMF, left) and formation of its microbial metabolite mycophenolic acid (MPA, right) under anaerobic conditions over time. Individual gut microbial communities are represented by color-coded lines. Lines and shaded areas depict the mean and s.e. of n=4 assay replicates, respectively. **(B)** Metabolism of MMF and formation of MPA over time by individual bacterial species under anaerobic conditions. *B.uniformis* used for MMF-metabolizing enzyme identification is depicted by a blue line, while the other 44 species are shown in gray. Lines and shaded areas depict the mean and s.e. of n=4 assay replicates, respectively. **(C)** Scheme for generation of an assayed gain-of-function (GoF) library and insert size distribution of the *B. uniformis* library based on 23 randomly picked clones. **(D)** Mapping of active insert sequences to the *B. uniformis* genome. **(E)** MMF-metabolizing activity of *B. uniformis* wild-type, BACUNI_RS05305 knockout and BACUNI_RS05305-complemented strains at different expression levels. Promoter strengths are P1E6 > P4E5 > P2E5 > P5E4 > P1E4 > P2E3 (*27*). Lines and shaded areas depict the mean and s.e. of n=4 assay replicates, respectively.

### Subhead 6: Using microbial signatures to explain microbial drug metabolism

We next sought to understand if community composition assessed by shotgun metagenomic analysis is predictive for microbiota drug metabolism. To this end, we trained random forest models to predict drug degradation (as continuous AUC values) from microbial species and gene abundance profiles of the *ex vivo* communities for which drug degradation was measured. Model performance was evaluated in leave-two-samples-out cross validation (treating anaerobic and microaerobic measurements from the same stool sample as different data points that were kept together either in the training or test subset). We contrasted these data-driven predictions with the ones based on selected species and genes that could be experimentally linked to specific drug metabolism in this or previous studies (Table S18) (*15, 19*). The analysis revealed pronounced differences between drugs and feature sets in how well drug metabolism could be predicted. Metabolization of the structurally similar drugs tacrolimus, everolimus and sirolimus could generally be predicted well (Spearman correlation between predicted and measured AUC between 0.64 and 0.68) using species abundance profiles. However, taking only those species as features that we identified as active metabolizers resulted in much poorer predictions not distinguishable from null models built with permuted labels (Spearman’s rho between 0.04 and 0.14, Fig. 6A, B). In contrast, when using genes with reported activity against tacrolimus, we observed much better predictions (achieving similar correlations as the species models) compared to purely data-driven models based on a large set of microbial gene abundances as features. Of note, given the similar metabolism of tacrolimus, everolimus, and sirolimus, the genes known to influence tacrolimus metabolism predicted the metabolism of sirolimus and everolimus similarly well. Feature importance analysis revealed that the models based on overall species abundance profiles for tacrolimus, everolimus and sirolimus all relied on a relatively small and consistent set of species as important predictors (Fig. S6). Predicting gut microbial metabolism of other drugs, most notably methylprednisolone and MMF, was found to be much more challenging. Only MMF predictions based on a large set of microbial gene abundances as features were slightly better than randomized null models (Spearman’s rho between -0.26 and 0.31) and neither species containing active strains, a larger set of abundant species, nor genes similar to the characterized MMF-metabolizing gene described above showed significant predictive capacity. To better predict MMF community metabolism, we first identified putative homologs (50% sequence identity, 50% coverage) of the MMF-metabolizing esterase that we experimentally identified in the genomes of our tested species. While we detected similar sequences even in species only distantly related to *B. uniformis*, the presence of such homologs only poorly predicted measured MMF degradation in these species (Fig. 6E, F). Motivated by this result and our observation that the closely related *B. uniformis* homolog BACUNI_RS05310 (with 85% sequence similarity) lacked a comparable metabolic function, we performed a structural analysis of BACUNI_RS05305 and BACUNI_RS05310 using AlphaFold2. We identified key differences in the substrate binding site, notably the substitution of methionine with alanine at position 191 within the oxyanion hole, alongside alterations in two arginine residues (Fig. 6D). These changes may affect the surface polarity and Van der Waals forces near the binding site, potentially impacting substrate specificity due to the role of the oxyanion hole in stabilizing the transition state during catalysis. Extending this structural analysis to the identified homologs across the tested species tended to better explain inactive species in which a sequence homologue had been detected, but was not sufficient to provide an explanation for all species with measured activity against MMF (Fig. 6E, F; Table S19). However, larger datasets (additional species and *ex vivo* communities with measured activity against MMF) will be needed to validate this approach. In summary, our study elucidates the complexities of predicting microbial metabolism of immunosuppressants, driven by the diverse genetics of microbial species, enzyme structures, and the chemical properties of drugs. While species-level information may be sufficient for some drugs, a deeper understanding of the responsible bacterial genes and structural details of enzyme-substrate interactions are crucial for others.

**Fig. 6.**
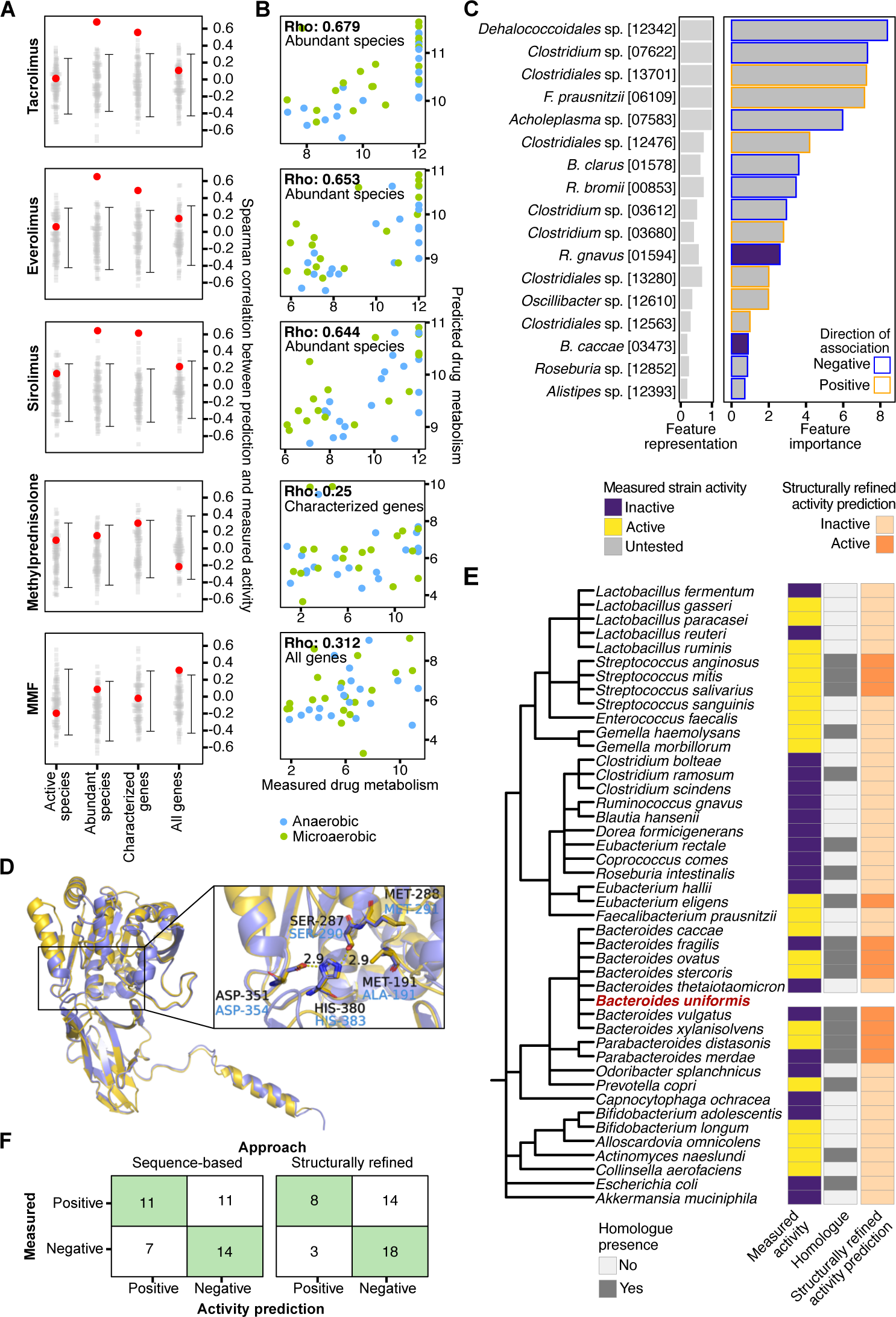
Predicting metabolic activity of microbial communities. **(A)** Random Forest cross-validation performances for four different models predicting community metabolism based on different sets of taxon- and gene-abundance features (see text) are denoted by red dots. These regression models predicted metabolic activity as quantified by the area under the drug concentration curve over time (AUC capped at 12) and were evaluated in terms of correlation between predicted and measured activity (see (B)). Gray dots indicate performance estimates of 100 null models trained with permuted labels; 90% confidence intervals are depicted by vertical lines with 5th and 95th percentile marked by horizontal ticks. **(B)** Scatter plots of measured versus predicted drug metabolism shown for the best-performing feature set evaluated in (A) for each drug (Spearman correlation denoted by rho). **(C)** Feature Importance (quantified by increase in mean squared error upon feature permutation, calculated on out-of-bag samples) of the most predictive taxa in the ‘abundant species’ model for tacrolimus. Feature representation corresponds to the fraction of cross-validated models that contain a given feature in their top feature set. Only features contained in at least 20% of models are shown. Directionality of association was inferred from the sign of the Spearman correlation between feature and response variable (metabolic activity). Positive associations (orange outline) indicate that a species is more abundant in communities with high metabolic activity. **(D)** Superposition of Alphafold2 predicted models for BACUNI_RS05305 (yellow, active enzyme) and BACUNI_RS05310 (blue, inactive enzyme) showing high structural similarity. Both paralogs are composed of a catalytic ‘/(-hydrolase domain and an immunoglobulin-like domain with an N-terminal signal peptide indicating secretion. The enlarged view shows the binding site with residues of the catalytic triad (Ser-His-Asp) and the oxyanion hole with hydrogen bonding distances of the catalytic triad given in Angstrom. Met191 in the active paralog is mutated to alanine in the inactive one, likely affecting (mycophenolate mofetil) MMF binding. **(E)** Taxonomy of the 44 species, for which MMF metabolism was measured anaerobically (first column), alongside detection of homologs to BACUNI_RS05305 (from *B. uniformis*, highlighted in red) inferred via sequence similarity search (second column), or via manual scoring of structural features of the active sites (see text) in these homologues (third column). **(F)** Contingency tables for bacterial MMF metabolism of 43 species (excluding *B. uniformis*) comparing activity prediction derived purely from sequence similarity-based homologue detection or additional structural refinement against measured activity.

## DISCUSSION

Microbial drug metabolism by single gut bacterial isolates has been well-documented for specific immunosuppressive drugs (*13, 16, 19*). In this study, we expanded this knowledge by systematically screening a broad range of immunosuppressive drugs of clinical relevance against a diverse array of human-derived gut microbial communities and bacterial isolates. Our findings revealed that nearly all immunosuppressive drugs undergo microbial biotransformation with marked interindividual and drug-specific variation. Intriguingly, we did not find universally strong or weak metabolizing communities, but rather that the ability of a given microbial community to metabolize one specific immunosuppressive drug at best weakly predicts its efficiency in metabolizing others (Fig. S7). This underscores the need to investigate gut microbial metabolism for each drug individually. Understanding and predicting microbial metabolism of immunosuppressive drugs in clinical settings will additionally be complicated by the fact that KT patients frequently receive a combination of tacrolimus, MMF, and methylprednisolone as standard first-line therapy.

Building on prior research that highlighted the influence of microbial metabolism on the efficacy and toxicity of immunosuppressive drugs (*20, 21*), our study further elucidates how gut microbiota can modify drug responses both locally and systemically. We demonstrated that bacteria can activate the prodrug MMF into its active form, mirroring the host’s prodrug activation. However, this bacterial activation may contribute to the gastrointestinal toxicity of MMF, as the local presence of the active metabolite MPA in the gut is linked to gastrointestinal adverse effects (*21*). On the other hand, the conversion of methylprednisolone into a metabolite with enhanced epithelial transport properties exemplifies the potential systemic impact of microbial drug metabolism. This is consistent with other studies demonstrating the systemic presence of microbial metabolites, such as the inactivated tacrolimus metabolite described above, in the bloodstream (*22*).

Our study of MMF, which pinpointed specific bacterial species and enzymes that metabolize this drug, contributes to expanding the mechanistic understanding of drug-microbiome interactions. Moreover, we extended previous observations on the bacterial inactivation of tacrolimus (*13*) by identifying related drug metabolites of sirolimus and everolimus, indicating a similar microbial inactivation process.

The challenges we faced in predicting microbial drug metabolism *in vitro* and *ex vivo* illustrate that predicting gut microbial metabolism of immunosuppressants in the clinical setting may be similarly complex. Additionally, our results suggest that the difficulty of predicting gut microbial drug metabolism may vary substantially among different immunosuppressants. While some compounds may only require taxonomic information, for others mechanistic information on metabolizing enzymes may be necessary. Moreover, the experimental identification of one metabolizing enzyme, even in an abundant gut commensal, may not be sufficient for a community-level understanding of drug metabolism, as suggested by inconsistencies between experimentally and model-identified key species. Such inconsistencies may however also arise from technical limitations inherent in both approaches (such as small sample size, Fig. 6C). Specifically, our work revealed limitations of using sequence similarity alone to predict community or species activity. While structural enzyme features may be used to refine homology searches aimed at quantifying the genomic abundance of similar drug-metabolizing genes in the community, the small size of our data set precludes a conclusive evaluation and rather highlights the challenges of predicting community metabolism. This is especially true for prevalent enzyme families, such as hydrolases, which are difficult to computationally quantify comprehensively in the microbiome as demonstrated with MMF.

In summary, our work adds to the increasing evidence for the influence of the gut microbiome on the metabolism of immunosuppressive drugs, but also highlights the difficulties in clinical translation of this knowledge. Nevertheless, by integrating our *in vitro* and *ex vivo* data with data from clinical studies, we may facilitate approaches towards personalized immunosuppression, recognizing the significant, but previously underappreciated role of the gut microbiome in individual variations in drug metabolism.

## MATERIALS AND METHODS

### Chemicals

Tested drugs were purchased from Sigma Aldrich, Santa Cruz Biotechnology or TCI as listed in Table S20. LC–MS-grade solvents were purchased from Guyer and other chemicals from Sigma Aldrich.

### Human fecal material

The study was approved by the Ethics Committee of the Medical Faculty of Heidelberg University (Study-ID: S-374/2021) and EMBL’s Bioethics Internal Advisory Board. It was performed in accordance with the World Medical Association Declaration of Helsinki Ethical Principles in the currently valid version. All samples were stored in the Zimmermann laboratory, using identifier numbers that are not associated with study volunteer names or other identifying information. Study volunteers were recruited at EMBL Heidelberg or the University Children’s Hospital Heidelberg. Table S1 gives an overview of all collected information of study volunteers. After collection samples were stored at 4°C and processed within < 18 hours under anaerobic conditions. Processing included the 1:1 addition of 40% glycerol in PBS followed by homogenization using the IKA ULTRA-TURRAX blender (VWR, Cat.Nr. 4312897). Samples were then aliquoted into 1 ml cryovials and stored at -80°C until further analysis.

### Microbial culture conditions

#### Anaerobic culture conditions

Anaerobic cultivation was carried out as previously described (*7*), using an anaerobic chamber (Coy Laboratory Products) with a gas mix of 20% CO2, 10% H2, and 70% N2. Solid cultures were grown on Brain-heart infusion (BHI; Becton Dickinson) agar with 10% horse blood. Liquid cultures of individual strains and fecal communities were grown in gut microbiota medium (GMM) (*23*).

#### Aerobic culture conditions

For molecular cloning, *E. coli* strains were aerobically cultured at 37°C in LB medium and on LB agar plates with kanamycin (50µg/ml), following previously established protocols (*7*).

### Mass spectrometry analysis of drugs and metabolites

#### Extraction of liquid samples

Liquid samples were processed for LC–MS analysis by organic solvent extraction as previously described (*7*). In brief, a mixture of acetonitrile:methanol (1:1) was introduced at -20°C following the addition of an internal standard mix, which included sulfamethoxazole, caffeine, ipriflavone, warfarin, and lisinopril, each at a final concentration of 80 nM. For samples derived from transwell assays, the internal standard mix excluded warfarin and lisinopril.

#### LC–MS analysis

Analysis was performed utilizing reversed-phase chromatography on an InfinityLab Poroshell HPH-C18 column (2.1×100 mm, 1.9 µm) with an Agilent 1200 Infinity UHPLC system. Mobile phases A (water with 0.1% formic acid) and B (methanol with 0.1% formic acid) were used, and the column was maintained at 45°C. A 5 µl sample was injected with an initial 100% phase A at a flow rate of 0.4 ml/min, followed by a linear gradient to 95% phase B over 5.5 minutes at the same flow rate. The qTOF (Agilent 6546) was operated in positive scanning mode (50–1,000 m/z), with source settings of 3,500 V for VCap, 2,000 V for nozzle voltage, a gas temperature of 225°C, a drying gas flow of 13 l/min, nebulizer pressure at 20 psig, and sheath gas settings of 225°C temperature and 12 l/min flow.

### Analysis of drug metabolism screen

#### Identification of metabolized drugs

A combined approach using both Fold Change (FC) and Area Under the Curve (AUC) was used to determine whether a drug was degraded. Fold change was used as reported previously (*7*). In brief, for each community, drug fold changes were calculated between each time point and 0 h in the 4 pools that contained a specific drug, and between these drug-containing pools and the non-drug controls. Statistical significance of the drug intensity differences was assessed with One-Way Permutation Test based on 9999 Monte-Carlo resampling (oneway_test function in R from the coin package) (***24***), and P values were FDR-corrected for multiple hypotheses testing using the Benjamini-Hochberg procedure (p.adjust function in R with method parameter set to “fdr”). To account for fast drug metabolism (within seconds after exposure), fold changes to control at time point 0 were used for drug and community/strain combinations for which (i) log2(fold change to control at t = 0) < −2; (ii) FDR-corrected P value (fold change to control at t = 0) < 0.05; and (iii) log2(fold change to control at t = 0) < log2(fold change at t = 12 h to t = 0). To account for variability in drug measurements, for each drug an adaptive fold-change threshold was calculated as either (i) (25%) or (ii) (mean + 2 s.d. of the fold changes, for which log2(fold change at t = 12 h to t = 0 h) > 0, to account for measurement noise), whichever was greater. Area Under the Curve was used as follows: AUC was computed using the R DescTools::AUC function on normalized and scaled areas for each of the sample, drug and pool combination. To account for variability in drug measurements, for each drug an adaptive AUC threshold was calculated as either (i) the average AUC computed on the control samples for the same drug, diminished of the desired percentage of metabolism (here, 25%), or (ii) the average AUC computed on the control samples for the same drug, diminished of two times their standard deviation whichever was smaller. A sample was reported as metabolized if its median AUC was less than the AUC threshold. Once both FC and AUC were used to assess drug degradation, if they returned two different results for the same drug-sample combination (*i.e.*, one identified it as degrading, the other as non-degrading), the following approach was used: (i) if AUC was claiming the sample as non-degrading the drug, then the FC result is kept only if either it is a quickly degrading drug or the fold change its double the requested FC; (ii) if AUC is claiming the sample as degrading the drug, then the AUC result is kept only if the corresponding FC is above 1 due to measurement noise at the 12h time point.

#### Identification of drug metabolites

We identified drug metabolite formation using linear mixed effects models predicting metabolite intensity using time and a random intercept for replicates. We assigned buildup if Benjamini-Hochberg corrected P value was < 0.1. To ensure the accuracy of metabolite intensity measurements, we adjusted for background noise by subtracting the baseline signal from the measured area before fitting the models.

### Preparing gain-of-function libraries, strain pooling and hit validation

Plasmids and primers are listed in Table S21 and S22.

#### Preparing gain-of-function library

Heterologous expression libraries were prepared as previously described (*7*). In brief, genomic DNA was extracted from overnight cultures of the source bacterial strain *B. uniformis* (*25*). DNA was fragmented into 2–8 kb pieces using focused ultrasonication (Covaris E220 with miniTUBE red). DNA Fragments were subsequently cloned into a PCR-linearized pZE21 expression vector (primers 1 and 2) through blunt-ended ligation (Thermo Fisher Rapid DNA Ligation kit). Ligated products were run on a 0.5% agarose gel to isolate fragments between 5 and 10 kb, which were then purified with a gel extraction kit (Qiagen). Ligation products were transformed into *E. cloni* 10G Elite competent cells (Lucigen) via electroporation. Colonies that grew overnight were selected and arrayed in a 384-well plate containing LB medium with kanamycin, assisted by a colony-picking robot (Singer Rotor). Inoculated plates were incubated overnight at 37°C, then duplicated onto LB agar plates with kanamycin and stored at -70°C for preservation.

#### Gain-of-function screen

Screening of the gain-of-function libraries was carried out as previously described (*7*). In brief, we first pooled all 384 colonies from a single library plate and resuspended them in diluted GMM containing MMF (5µM). The suspension was incubated anaerobically at 37°C and samples were collected at intervals of 0, 1, 2, 4, 6, 8, 12, and 24 hours during incubation. Pools showing the ability to metabolize MMF were then transferred to a new 384-well plate and cultured aerobically for 12 hours at 37°C. We identified three active plates and we created pools of each row and column for further investigation. Active clones that metabolized MMF were isolated, and four individual colonies from each were re-evaluated for their MMF-metabolizing capability. Once confirmed, these clones underwent Sanger sequencing to identify the genetic inserts using primers 3 and 4.

#### Hit validation by targeted gene expression in E. coli

To confirm the activity of the identified genes, their sequences were amplified using PCR with primers 5 and 6, cloned into the pZE21 expression vector via Gibson cloning (NEBuilder HiFi DNA Assembly Kit from NEB), and then transformed into *E. cloni* 10G Elite cells by electroporation. The resulting bacterial strains were further tested for MMF metabolization.

### Construction of *B. uniformis* targeted mutant and complementation strains

Bacterial strains, plasmids, and primers are listed in Tables S6, 21 and S22. Gene deletions and complementations were carried out as previously described (*26*). The BACUNI_RS05305 gene was deleted from *B. uniformis* (DSM6597) using a pLGB13 suicide plasmid. The plasmid was constructed using primer numbers 7-14. For introducing the plasmid into *B. uniformis*, conjugation with *E. coli* S17-pir was utilized. *E. coli* S17-pir containing plasmid pLGB13_MMF_KO_BACUNIRS05305 was cultured aerobically in liquid LB medium supplemented with ampicillin (100µg/ml), while *B. uniformis* was grown anaerobically in BHI medium enriched with hematin (0.5 mg/mL) and menadione (0.1 mg/mL). Following growth, both bacterial cultures were pelleted by centrifugation, resuspended, and mixed in 50 µL of BHI medium supplemented with hematin and menadione. This mixture was then spotted onto a BHI-blood agar plate and incubated aerobically at 37°C for 12-24 hours. After incubation, the bacterial culture was transferred onto BHI agar plates containing erythromycin (5 µg/mL) and gentamicin (200 µg/mL) to select for transconjugant *B. uniformis* clones. After 2 days of anaerobic incubation, colonies were picked and further screened for second recombination events on BHI plates containing anhydrotetracycline (100 ng/mL) for counterselection. Verification of scare-less gene knock outs was performed using PCR with primers 14 and 15. Gene complementation at different expression levels using engineered phage promotors were performed with primer numbers 16-19 as described before (*7, 27*).

### Epithelial transport assay

Caco-2 cells were obtained from Sigma (Cat no. 86010202) at passage 9. Cells used in this study were not passaged more than 20 times. Transwell inserts were obtained from Millicell, and were used in combination with 24-well plates (Thermo Fisher). The Caco-2 cells were grown in T75 flasks at 37°C in an incubator with an atmosphere of 5% CO2. The cells were grown using Dulbecco’s modified Eagle medium (DMEM) (Gibco, 41965-039) with 4.5 g/liter D-glucose, 584 mg/liter L-glutamine, 1% penicillin/streptomycin, 1% amphotericin B, and 10% fetal bovine serum. The medium in the flasks was changed every 3 days. At 90% confluence, the cells were passaged using incubation with 0.5% trypsin for 5 minutes at 37°C after washing the cells with PBS. Cells were then seeded into the transwell inserts (pore size 0.4 μm) at 10,000 cells/well (Merck). The transwells with the Caco-2 cells were used only after >21 days post-seeding to allow for the establishment of a stable monolayer. During this post-seeding period, the media was changed every 3 days. For each assay, the TEER of each monolayer was measured, to ensure the stability and permeability of each monolayer. Transwells were only used when the TEER measurement was above 2,083 Ohm/cm2 per well. The transwell inserts with the Caco-2 monolayer were washed 2 times with Hank’s Buffered Salt Solution (HBSS), before adding the sterile supernatant from the bacterial strains, or the single compounds in solution to the apical or basolateral side. Small volumes of both the apical and basolateral side (10 uL) were taken every 30 minutes and immediately stored at -70°C until metabolite extraction. Time points were taken for 5 hours, whereafter the cells were washed 3 times with HBSS and media was added to the cells, on the apical side containing a viability dye (Promega, G8080). After an overnight incubation, the apical media was transferred to a separate 96-well plate and fluorescent signal was measured. Quantification of the compound concentrations in the epithelial transport assays was based on a dilution series of chemical standards spanning 0.001 to 10 μM. The effective permeability coefficients from the receiver compartments were calculated as described previously (*18*). In short, the following formula was used: 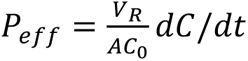. Here, *V_R_* is the volume of the receiver compartment, *A* the transwell surface area, *C_0_* the initial concentration of the solute in the receiver compartment, and *dC/dt* the slope of the first 240 minutes of the concentration of the solute in the receiver compartment. Statistical analysis and plotting were performed in RStudio Version 4.4.2.

### Metagenomic sequencing analysis

#### Sample preparation and sequencing

Fecal DNA was extracted from a biomass pellet of 1.5 ml of microbial community cultures using the ZymoBIOMICS DNA Miniprep Kit (D4300). Library preparation and sequencing were performed at the EMBL Genomics Core Facility at EMBL Heidelberg. The NEBNext Ultra II DNA kit was used for the preparation of the metagenomic sequencing libraries. Sequencing was performed on an Illumina NextSeq2000 instrument with a P3 flow-cell to target depth of 40 million reads per sample.

#### Taxonomic profiling

Raw shotgun metagenomic reads were subjected to the following read cleaning protocol with bbduk (bbmap-version 38.93): 1) low quality trimming on either side (qtrim = rl, trimq = 3), 2) discarding of low-quality reads (maq = 25), 3) adapter removal (ktrim = r, k = 23, mink = 11, hadis = 1, tpe = true, tbo = true; against the bbduk default adapter library) and 4) length filtering (ml = 45). The cleaned reads were screened for host contamination using kraken2 (version 2.1.2) against the human hg38 reference genome with ribosomal sequences masked (SILVA 138). Microbial abundance profiles were then generated using mOTUs3 with default settings (*28*).

#### Functional profiling

Prior to functional profiling, raw shotgun metagenomics reads were subjected to the same read cleaning protocol as described in the section ‘Taxonomic profiling’. Preprocessed reads were then mapped to the human gut specific Global Microbial Gene Catalogue (GMGC) (*29*) using BWA-MEM (0.7.17) with default parameters. Here, a reduced GMGC catalog excluding very rare genes was used (described in (*30*)). Alignments were filtered to >45bp alignment length and >97% sequence identity and profiled with gffquant version v2.9.1 (https://github.com/cschu/gff_quantifier). Reads aligning ambiguously to multiple genes contributed proportionally.

### Predicting community microbial activity from gene- and taxon abundances

In order to predict community-wide metabolic activity, we built Random Forest models to infer observed drug degradation (measured via AUC, defined above) using either community mOTU-level taxonomic profiles or gene-level functional profiles. For model building, we used 38 metagenomic samples (19 anaerobic and 19 microaerobic). Models were trained and evaluated in a leave-two-samples-out manner where samples from the same donor were always kept together in the training or test set. We evaluated four different feature sets defined as follows: ‘Active species’ correspond to those mOTUs that contain a metabolically active strain. Mapping between strains and mOTUs was achieved by extracting mOTUs marker genes from strain genomes and subsequent assignment to mOTUs (using functionality provided with mOTUs3 https://github.com/motu-tool/mOTUs-extender). ‘Abundant species’ features were derived from relative abundances of the 345 mOTUs with prevalence > 20% and maximum abundance across samples of > 0.1%. ‘Characterized genes’ correspond to the GMGC clusters that are similar to an enzyme with established metabolic activity (Table S18), where similarity was defined as gene sequences in the GMGC being at least 40% similar over at least 70% of their length, determined using DIAMOND (*31*). ‘All genes’ corresponds to all ∼9M GMGC genes found in the *ex vivo* communities. To reduce this feature set to a manageable size before Random Forest training, we calculated feature-wise correlations to the label on each training set and retained only the best-correlated 1000 features as input to the Random Forest. For all predictions, input features were standardized (to mean 0 and standard deviation 1) before training the model.

In order to define the set of most important features for all evaluated models (shown in Fig. 6 and S6), we trained models by recursive feature addition, *i.e.*, successively adding the most predictive features (computed via out-of-bag-estimated % increase in MSE upon feature permutation using the RandomForest function in R) until the model based on this small set of top features achieved a test performance larger than 90% of the full model. This was conducted for each leave-two-samples-out fold individually, which resulted in a fold-specific set of top features, across which information was aggregated to display feature importance (shown in Fig. 6 and S6).

### Predicting drug-metabolizing capacity of bacterial strains

In order to facilitate prediction of MMF-metabolic activity of gut communities, homologs of BACUNI_RS05305 were identified in the genomes of 43 bacterial strains using a BLAST (*32*) search using 50% sequence identity and 50% query coverage as cutoffs. We excluded *B. uniformis* from this search as the query enzyme originated from this organism. In total, 32 putative homologs were detected in 18 strains. To structurally investigate the active sites of these homologs, Alphafold2 v2.3.2 (*33, 34*) was used with default parameters to predict five models for each of the putative homologs. The highest confidence model was relaxed and then used for further analysis. Conservation of the catalytic triad and the oxyanion hole residues was assessed through a multiple sequence alignment of all 32 homologs using MUSCLE v5 (*35*). After alignment on catalytic residues, accessibility and polarity of the binding site was assessed by visual inspection. We then used structural features around the catalytic site to obtain structure-informed activity predictions. A fully accessible substrate binding site was considered necessary to predict an enzyme to be active. Additionally, either a fully apolar binding site (no charged, surface-exposed residues within 5A) or a binding site featuring both charged residues and large apolar residues composing the oxyanion hole at position 191 was deemed necessary for activity against MMF. Finally, we predicted a strain to be active against MMF, if a single putative homolog in this strain was predicted active. As a result, of the 32 detected homologs from 18 strains, 24 homologs from 11 strains were classified as active using this structurally refined prediction. Images of structures were generated using open source PyMOL (PyMOL Molecular Graphics System, Version 2.3.0, Schrödinger, LLC.)

## Acknowledgments

We thank the Zeller and the Zimmerman labs for stimulating discussions of our work and the EMBL GeneCore Facility for services and experimental advice.

## Author contributions

Conceptualization: MBA, DC, BT, GZ, MZ

Methodology: GZ, MZ

Investigation: MBA, NK, EM, RH, MG, AB

Visualization: MBA, NK, AB, EM, RH

Funding acquisition: MBA, BT, GZ, MZ

Project administration: MBA

Supervision: BT, GZ, MZ

Writing – original draft: MBA

Writing – review & editing: MBA, NK, AB, EM, RH, MG, DC, BT, GZ, MZ

## Data and materials availability

All data are available in the main text or the supplementary materials. Raw metagenomic sequencing data will be made available under the ENA project ID PRJEB74094. Taxonomic and functional profiles are available under the Zenodo accession ID 10895043. Raw metabolomics data will be made available in the MetaboLights repository, with accession number MTBLS9827. Code for analyzing data and generating figures are available on GitHub (https://github.com/zellerlab/gut-microbial-metabolism-of-immunosuppressants).

## References and Notes

1. A. Rana, A. Gruessner, V. G. Agopian, Z. Khalpey, I. B. Riaz, B. Kaplan, K. J. Halazun, R. W. Busuttil, R. W. G. Gruessner, Survival Benefit of Solid-Organ Transplant in the United States. JAMA Surg 150, 252 (2015).

2. M. Coemans, C. Süsal, B. Döhler, D. Anglicheau, M. Giral, O. Bestard, C. Legendre, M.-P. Emonds, D. Kuypers, G. Molenberghs, G. Verbeke, M. Naesens, Analyses of the short- and long-term graft survival after kidney transplantation in Europe between 1986 and 2015. Kidney International 94, 964–973 (2018).

3. S. Bergan, M. Brunet, D. A. Hesselink, K. L. Johnson-Davis, P. K. Kunicki, F. Lemaitre, P. Marquet, M. Molinaro, O. Noceti, S. Pattanaik, T. Pawinski, C. Seger, M. Shipkova, J. J. Swen, T. van Gelder, R. Venkataramanan, E. Wieland, J.-B. Woillard, T. C. Zwart, M. J. Barten, K. Budde, M.-T. Dieterlen, L. Elens, V. Haufroid, S. Masuda, O. Millan, T. Mizuno, D. J. A. R. Moes, M. Oellerich, N. Picard, L. Salzmann, B. Tönshoff, R. H. N. van Schaik, N. T. Vethe, A. A. Vinks, P. Wallemacq, A. Åsberg, L. J. Langman, Personalized Therapy for Mycophenolate: Consensus Report by the International Association of Therapeutic Drug Monitoring and Clinical Toxicology. Therapeutic Drug Monitoring 43, 150–200 (2021).

4. M. Brunet, T. van Gelder, A. Åsberg, V. Haufroid, D. A. Hesselink, L. Langman, F. Lemaitre, P. Marquet, C. Seger, M. Shipkova, A. Vinks, P. Wallemacq, E. Wieland, J. B. Woillard, M. J. Barten, K. Budde, H. Colom, M.-T. Dieterlen, L. Elens, K. L. Johnson-Davis, I. MacPhee, S. Masuda, B. S. Mathew, O. Millán, T. Mizuno, D.-J. A. R. Moes, C. Monchaud, O. Noceti, T. Pawinski, N. Picard, R. van Schaik, C. Sommerer, N. T. Vethe, B. de Winter, U. Christians, S. Bergan, Therapeutic Drug Monitoring of Tacrolimus-Personalized Therapy: Second Consensus Report. Ther Drug Monit 41, 47 (2019).

5. N. Shuker, T. van Gelder, D. A. Hesselink, Intra-patient variability in tacrolimus exposure: Causes, consequences for clinical management. Transplantation Reviews 29, 78–84 (2015).

6. L. M. Andrews, Y. Li, B. C. M. De Winter, Y.-Y. Shi, C. C. Baan, T. Van Gelder, D. A. Hesselink, Pharmacokinetic considerations related to therapeutic drug monitoring of tacrolimus in kidney transplant patients. Expert Opinion on Drug Metabolism & Toxicology 13, 1225–1236 (2017).

7. M. Zimmermann, M. Zimmermann-Kogadeeva, R. Wegmann, A. L. Goodman, Mapping human microbiome drug metabolism by gut bacteria and their genes. Nature 570, 462–467 (2019).

8. B. Javdan, J. G. Lopez, P. Chankhamjon, Y.-C. J. Lee, R. Hull, Q. Wu, X. Wang, S. Chatterjee, M. S. Donia, Personalized Mapping of Drug Metabolism by the Human Gut Microbiome. Cell (2020), doi:10.1016/j.cell.2020.05.001.

9. A. A. Verdegaal, A. L. Goodman, Integrating the gut microbiome and pharmacology. Sci. Transl. Med. 16, eadg8357 (2024).

10. M. Zimmermann, K. R. Patil, A. Typas, L. Maier, Towards a mechanistic understanding of reciprocal drug–microbiome interactions. Molecular Systems Biology 17, e10116 (2021).

11. K. A. Lee, Cross-cohort gut microbiome associations with immune checkpoint inhibitor response in advanced melanoma. Nature Medicine, 28.

12. M. R. Taylor, K. L. Flannigan, H. Rahim, A. Mohamud, I. A. Lewis, S. A. Hirota, S. C. Greenway, Vancomycin relieves mycophenolate mofetil–induced gastrointestinal toxicity by eliminating gut bacterial β-glucuronidase activity. Science Advances 5, eaax2358 (2019).

13. Y. Guo, C. M. Crnkovic, K.-J. Won, X. Yang, J. R. Lee, J. Orjala, H. Lee, H. Jeong, Commensal gut bacteria convert the immunosuppressant tacrolimus to less potent metabolites. Drug Metabolism and Disposition, dmd.118.084772 (2018).

14. G. P. Donaldson, S. M. Lee, S. K. Mazmanian, Gut biogeography of the bacterial microbiota. Nature Reviews Microbiology 14, 20–32 (2016).

15. S. Devendran, S. M. Mythen, J. M. Ridlon, The *desA* and *desB* genes from *Clostridium scindens* ATCC 35704 encode steroid-17,20-desmolase. Journal of Lipid Research 59, 1005– 1014 (2018).

16. M. P. M. Letertre, N. Munjoma, K. Wolfer, A. Pechlivanis, J. A. K. McDonald, R. N. Hardwick, N. J. Cherrington, M. Coen, J. K. Nicholson, L. Hoyles, J. R. Swann, I. D. Wilson, A Two-Way Interaction between Methotrexate and the Gut Microbiota of Male Sprague–Dawley Rats. J. Proteome Res. 19, 3326–3339 (2020).

17. S. Lazarević, M. Đanic, H. Al-Salami, A. Mooranian, M. Mikov, Gut Microbiota Metabolism of Azathioprine: A New Hallmark for Personalized Drug-Targeted Therapy of Chronic Inflammatory Bowel Disease. Front. Pharmacol. 13, 879170 (2022).

18. A. R. Hilgers, R. A. Conradi, P. S. Burton, Caco-2 Cell Monolayers as a Model for Drug Transport Across the Intestinal Mucosa. Pharmaceutical Research 07, 902–910 (1990).

19. L. Qian, H. Ouyang, L. Gordils-Valentin, J. Hong, A. Jayaraman, X. Zhu, Identification of Gut Bacterial Enzymes for Keto-Reductive Metabolism of Xenobiotics. ACS Chem. Biol. 17, 1665–1671 (2022).

20. A. L. Degraeve, V. Haufroid, A. Loriot, L. Gatto, V. Andries, L. Vereecke, L. Elens, L. B. Bindels, Gut microbiome modulates tacrolimus pharmacokinetics through the transcriptional regulation of ABCB1. Microbiome 11, 138 (2023).

21. P. Li, R. Zhang, J. Zhou, P. Guo, Y. Liu, S. Shi, Vancomycin relieves tacrolimus-induced hyperglycemia by eliminating gut bacterial beta-glucuronidase enzyme activity. Gut Microbes 16, 2310277 (2024).

22. Y. Guo, H. Lee, E. Edusei, S. Albakry, H. Jeong, J. R. Lee, Blood Profiles of Gut Bacterial Tacrolimus Metabolite in Kidney Transplant Recipients., 2.

23. A. L. Goodman, G. Kallstrom, J. J. Faith, A. Reyes, A. Moore, G. Dantas, J. I. Gordon, Extensive personal human gut microbiota culture collections characterized and manipulated in gnotobiotic mice. Proc Natl Acad Sci U S A 108, 6252–6257 (2011).

24. T. Hothorn, K. Hornik, M. A. V. D. Wiel, A. Zeileis, Implementing a Class of Permutation Tests: The **coin** Package. J. Stat. Soft. 28 (2008), doi:10.18637/jss.v028.i08.

25. A. L. Goodman, M. Wu, J. I. Gordon, Identifying microbial fitness determinants by insertion sequencing using genome-wide transposon mutant libraries. Nat Protoc 6, 1969–1980 (2011).

26. L. García-Bayona, L. E. Comstock, K. P. Lemon, Ed. Streamlined Genetic Manipulation of Diverse Bacteroides and Parabacteroides Isolates from the Human Gut Microbiota. mBio 10, e01762–19 (2019).

27. W. R. Whitaker, E. S. Shepherd, J. L. Sonnenburg, Tunable Expression Tools Enable Single-Cell Strain Distinction in the Gut Microbiome. Cell 169, 538–546.e12 (2017).

28. H.-J. Ruscheweyh, A. Milanese, L. Paoli, N. Karcher, Q. Clayssen, M. I. Keller, J. Wirbel, P. Bork, D. R. Mende, G. Zeller, S. Sunagawa, Cultivation-independent genomes greatly expand taxonomic-profiling capabilities of mOTUs across various environments. Microbiome 10, 212 (2022).

29. L. P. Coelho, R. Alves, Á. R. Del Río, P. N. Myers, C. P. Cantalapiedra, J. Giner-Lamia, T. S. Schmidt, D. R. Mende, A. Orakov, I. Letunic, F. Hildebrand, T. Van Rossum, S. K. Forslund, S. Khedkar, O. M. Maistrenko, S. Pan, L. Jia, P. Ferretti, S. Sunagawa, X.-M. Zhao, H. B. Nielsen, J. Huerta-Cepas, P. Bork, Towards the biogeography of prokaryotic genes. Nature 601, 252–256 (2022).

30. Q. R. Ducarmon, N. Karcher, H. L. P. Tytgat, O. Delannoy-Bruno, S. Pekel, F. Springer, C. Schudoma, G. Zeller, Large-scale computational analyses of gut microbial CAZyme repertoires enabled by Cayman (bioRxiv 2024.01.08.574624, 2024; 10.1101/2024.01.08.574624).

31. B. Buchfink, C. Xie, D. H. Huson, Fast and sensitive protein alignment using DIAMOND. Nat Methods 12, 59–60 (2015).

32. C. Camacho, G. Coulouris, V. Avagyan, N. Ma, J. Papadopoulos, K. Bealer, T. L. Madden, BLAST+: architecture and applications. BMC Bioinformatics 10, 421 (2009).

33. M. Varadi, S. Anyango, M. Deshpande, S. Nair, C. Natassia, G. Yordanova, D. Yuan, O. Stroe, G. Wood, A. Laydon, A. Žídek, T. Green, K. Tunyasuvunakool, S. Petersen, J. Jumper, E. Clancy, R. Green, A. Vora, M. Lutfi, M. Figurnov, A. Cowie, N. Hobbs, P. Kohli, G. Kleywegt, E. Birney, D. Hassabis, S. Velankar, AlphaFold Protein Structure Database: massively expanding the structural coverage of protein-sequence space with high-accuracy models. Nucleic Acids Research 50, D439–D444 (2022).

34. J. Jumper, R. Evans, A. Pritzel, T. Green, M. Figurnov, O. Ronneberger, K. Tunyasuvunakool, R. Bates, A. Žídek, A. Potapenko, A. Bridgland, C. Meyer, S. A. A. Kohl, A. J. Ballard, A. Cowie, B. Romera-Paredes, S. Nikolov, R. Jain, J. Adler, T. Back, S. Petersen, D. Reiman, E. Clancy, M. Zielinski, M. Steinegger, M. Pacholska, T. Berghammer, S. Bodenstein, D. Silver, O. Vinyals, A. W. Senior, K. Kavukcuoglu, P. Kohli, D. Hassabis, Highly accurate protein structure prediction with AlphaFold. Nature 596, 583–589 (2021).

35. R. C. Edgar, Muscle5: High-accuracy alignment ensembles enable unbiased assessments of sequence homology and phylogeny. Nat Commun 13, 6968 (2022).

