## Supplementary material for "Unraveling interindividual differences and functional consequences of gut microbial metabolism of immunosuppressants": Suppl. figures 1-8


Maral Baghai Arassi *et al.*


**This PDF file includes:**

Figs. S1 to S8

Tables S1 to S22 (only headers, tables in separate file)

**Fig. S1.**

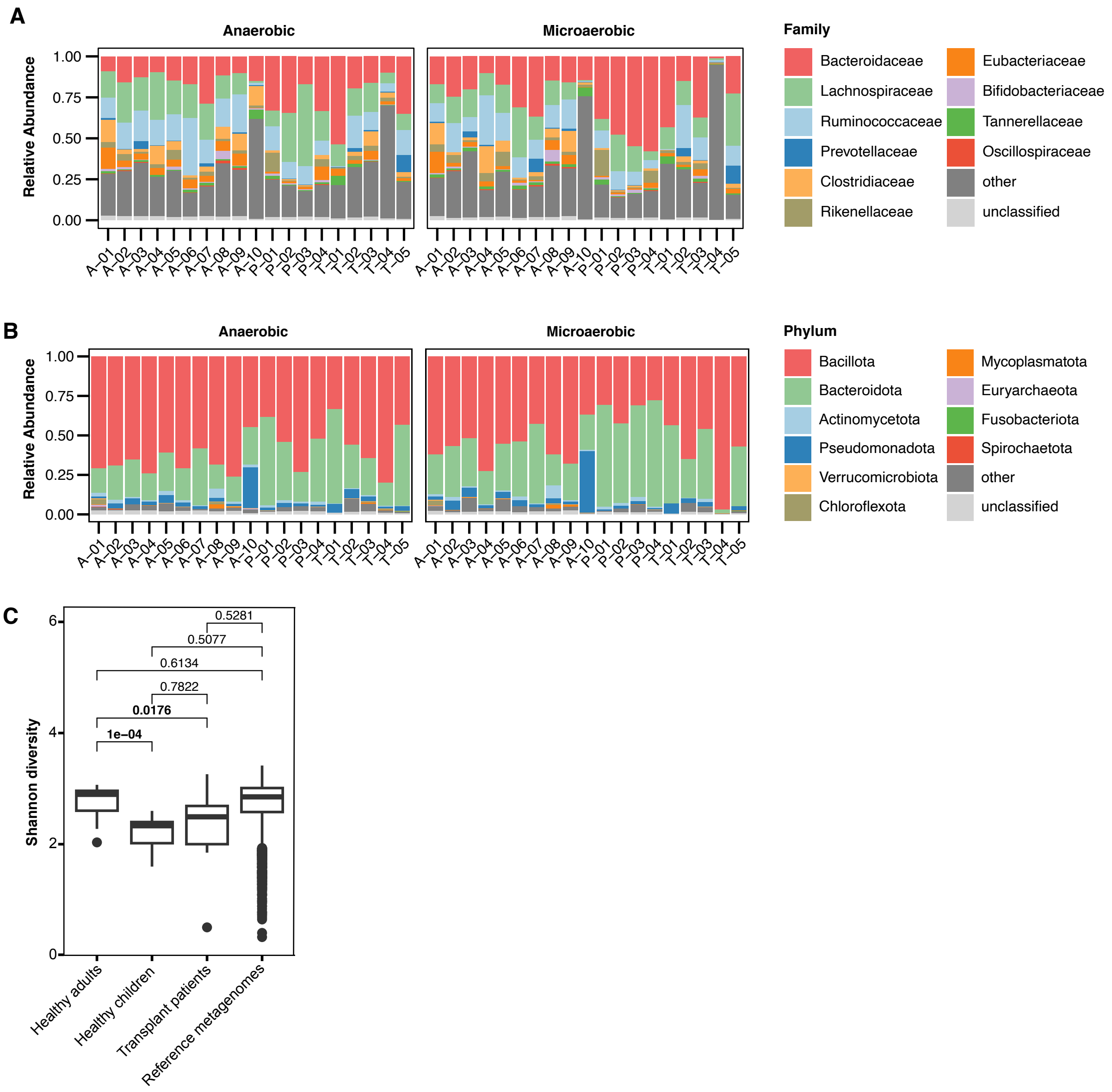


**Fig. S1. Bacterial relative abundances of gut microbial communities.**

Relative abundances of bacterial taxa in the 38 human-derived gut microbial communities on family **(A)** and phylum **(B)** level under anaerobic and microaerobic conditions. **(C)** Shannon diversity of human-derived gut microbial communities compared to the Shannon diversity in the collection of reference metagenomes.

**Fig. S2.**

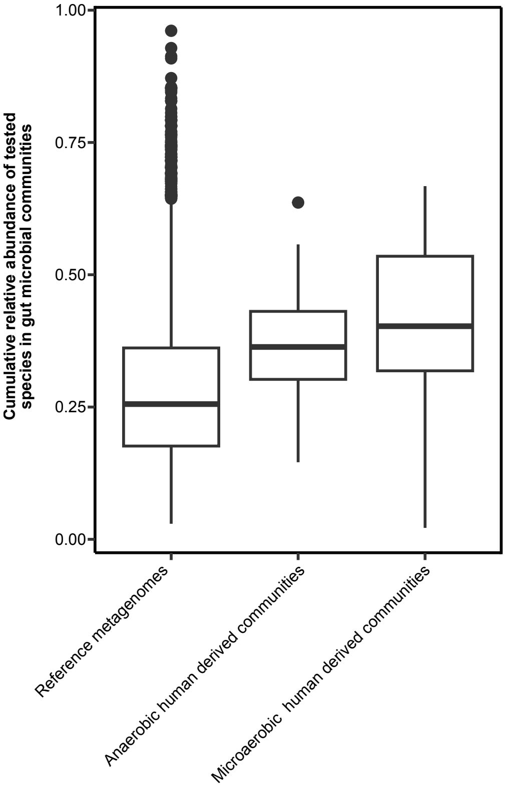


**Fig. S2. Cumulative relative abundance of tested species in gut microbial communities.**

Cumulative relative abundance of tested species in the 38 human-derived microbial communities under anaerobic and microaerobic conditions as well as in the reference metagenomes.

Fig. S3.

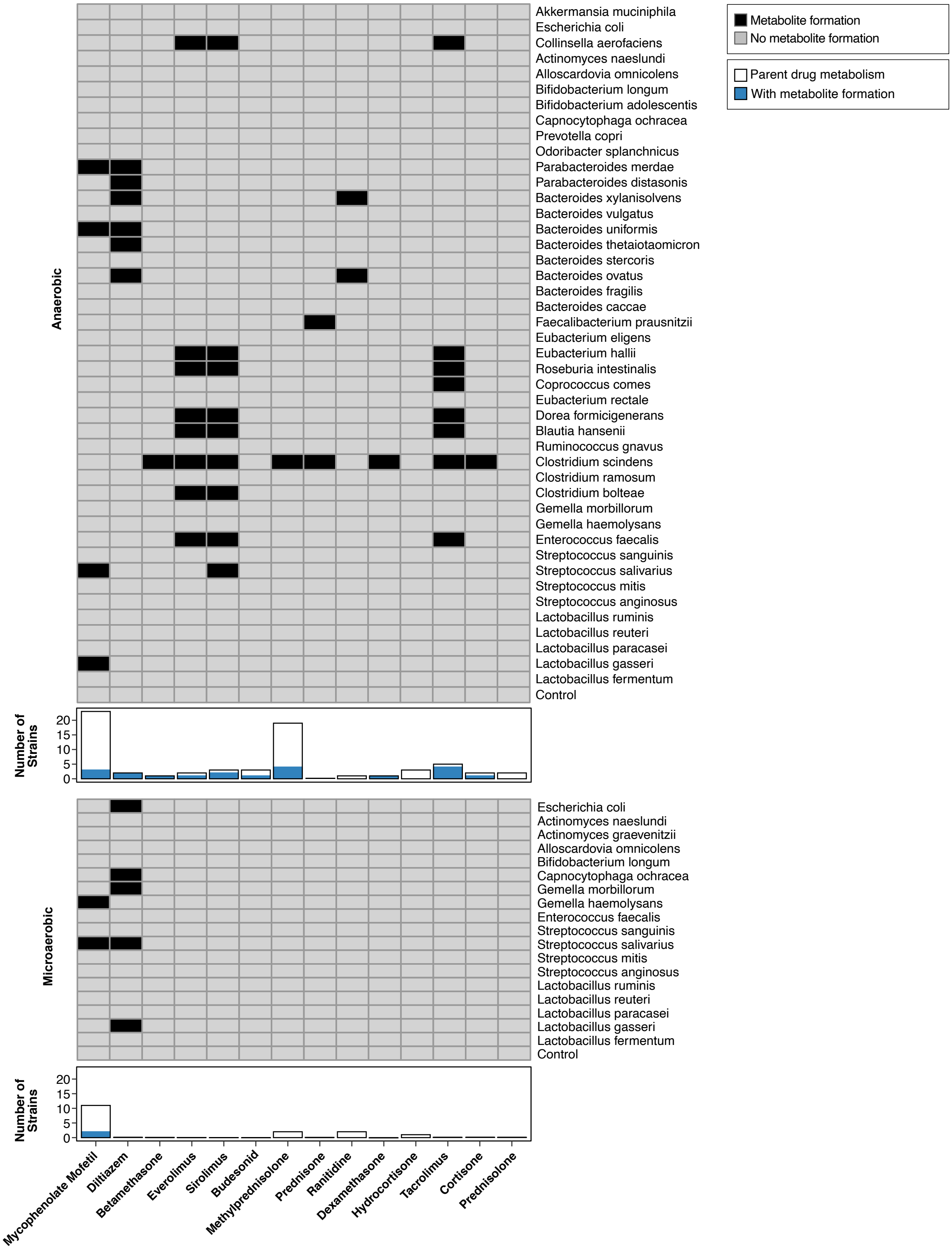


**Fig. S3. Identified drug metabolites from individual gut bacterial species.**

**(A)** Heat map illustrating microbial drug metabolite formation based on the parent drug. Barplots below the heat map depict the number of species metabolizing a given parent drug. Bar outlines below the heat map depict the number of species metabolizing a given parent drug. Blue filled bars represent the number of species where additional metabolite formation was observed. Drugs are sorted as in Fig. 3, while microbial species are sorted based on taxonomy.

**Fig. S4.**

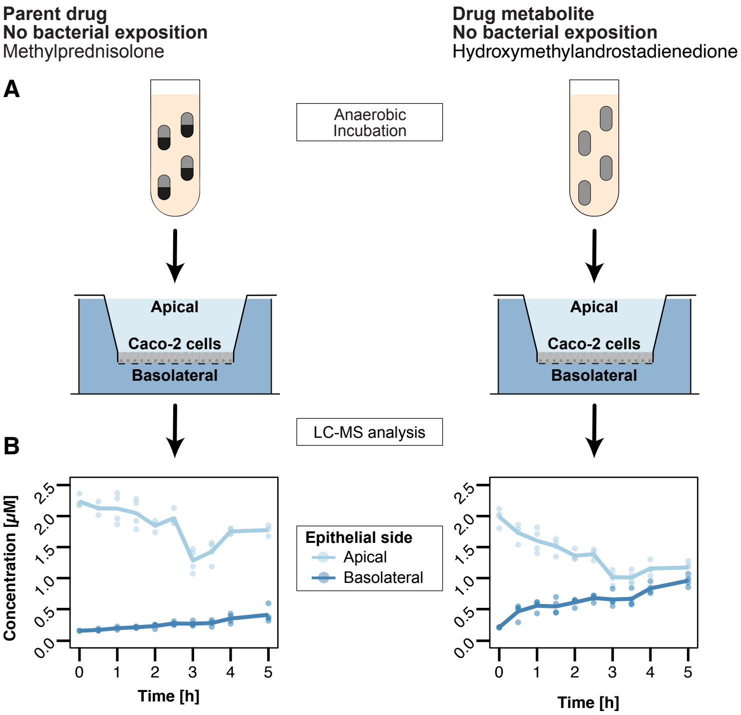


**Fig. S4. Epithelial transport characteristics of methylprednisolone and its microbial metabolite under sterile conditions.**

**(A)** Schematic of the transport assay **(B)** Line plot depicting the concentration over time of methylprednisolone (left) and its microbial metabolite hydroxymethylandrostadienedione (right) on the apical and basolateral side, respectively. Lines represent the mean of n=4 assay replicates while circles represent individual measurements.

**Fig. S5.**

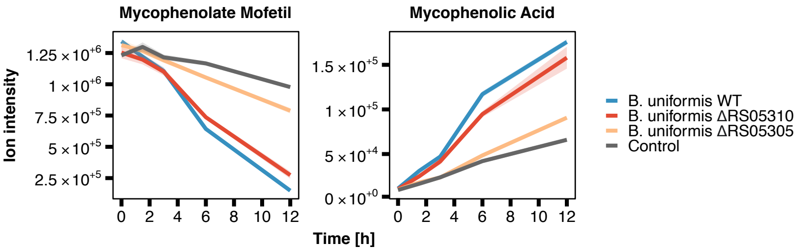


**Fig. S5. Comparison of the identified bacterial MMF-metabolizing gene product with an inactive homolog.**

MMF-metabolizing activity of *B. uniformis* wild-type, BACUNI_RS05305 knockout and BACUNI_RS05310 knockout. Lines and shaded areas depict the mean and s.e. of n=4 assay replicates, respectively.

**Fig. S6.**

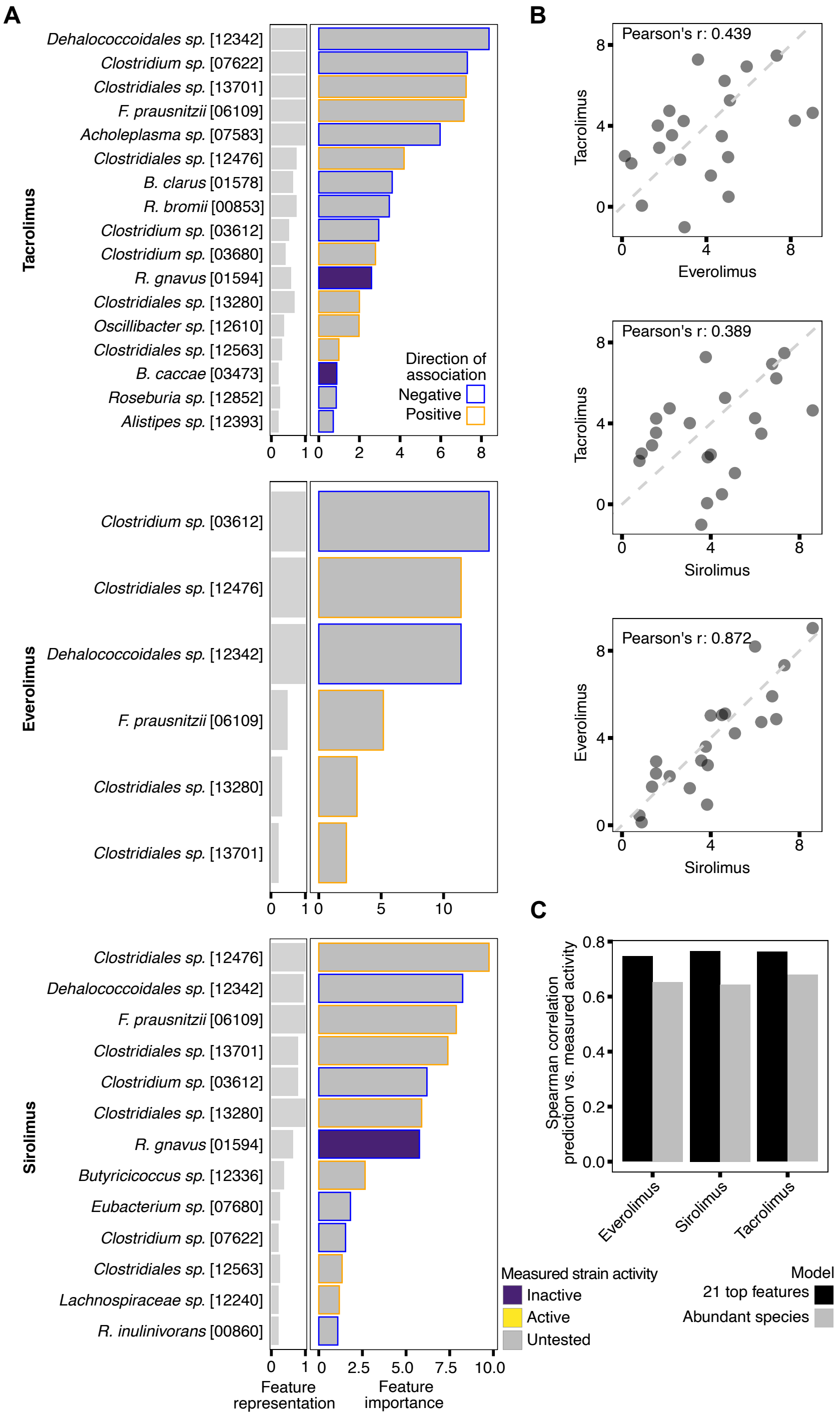


**Fig. S6. Comparison of models predicting tacrolimus, sirolimus and everolimus metabolic activity in communities.**

**(A)** Most important species for predicting metabolization of tacrolimus, sirolimus and everolimus in microbial communities. Computed analogously to Fig. 6C. **(B)** Pairwise scatter plots of feature importance values of models predicting tacrolimus, sirolimus and everolimus. Feature importance values were computed on models retrained on the superset of the features shown in (A), equalling a total of 21 distinct bacterial species. **(C)** Spearman correlations between measured and predicted drug AUC values for 'abundant species' models shown in Fig. 6 as well as for models trained on 21 features.

**Fig. S7.**

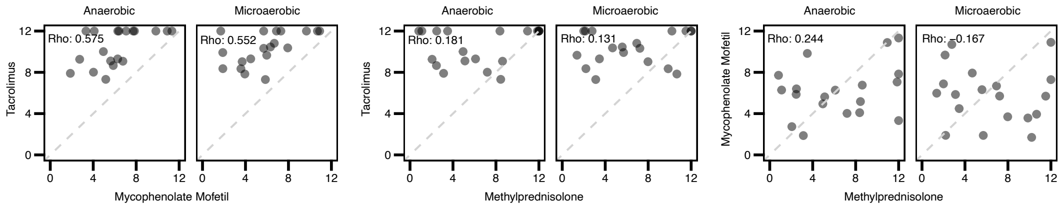


**Fig. S7. Correlation of microbial metabolism of tacrolimus, mycophenolate mofetil and methylprednisolone.**

Pairwise scatter plots of community metabolic activity (measured by area under the drug concentration curve, AUC) for tacrolimus, mycophenolate mofetil and methylprednisolone under anaerobic and microaerobic oxygen conditions. Rho indicates Spearman correlation.

**Fig. S8.**

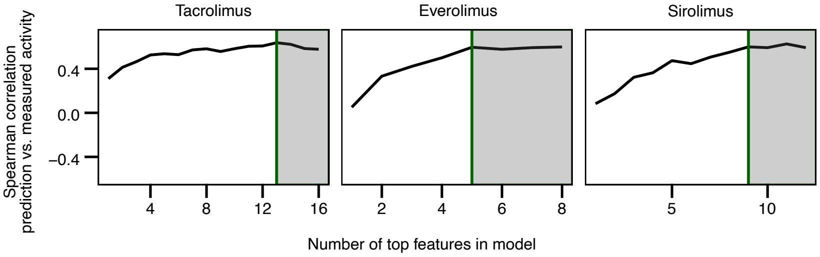


**Fig. S8. Model performance for tacrolimus, everolimus and sirolimus as a function of the number of features.**

Line plots relating number of top features to model performance for tacrolimus, everolimus and sirolimus. Green vertical line corresponds to the cutoff where the model is within 10% performance (measured by Spearman correlation between measured and predicted AUC) of the corresponding full model.

**Table S1**. Human stool donor information

**Table S2**. Reference metagenomes used in this study

**Table S3**. Metagenomics data: bacterial community composition anaerobic (overnight cultures)

**Table S4**. Metagenomics data: bacterial community composition microaerobic (overnight cultures)

**Table S5**. Drugs used in this study, name, formula, mass, chromatographic retention time

**Table S6**. Drug screen results (parent drugs), fold changes and p-values for human fecal community-drug interactions (anaerobic)

**Table S7**. Drug screen results (parent drugs), fold changes and p-values for human fecal community-drug interactions (microaerobic)

**Table S8**. Human gut bacteria tested for drug-metabolizing activity

**Table S9**. Drug screen results (parent drugs), fold changes and p-values for all single species-drug interactions (anaerobic)

**Table S10**. Drug screen results (parent drugs), fold changes and p-values for all single species-drug interactions (microaerobic)

**Table S11**. Metabolites used for suspect screening of untargeted metabolomics data

**Table S12**. Drug screen results (metabolites), fold changes and p-values for human fecal community-drug interactions (anaerobic)

**Table S13**. Drug screen results (metabolites), fold changes and p-values for human fecal community-drug interactions (microaerobic)

**Table S14**. Drug screen results (metabolites), fold changes and p-values for all single species-drug interactions (anaerobic)

**Table S15**. Drug screen results (metabolites), fold changes and p-values for all single species-drug interactions (microaerobic)

**Table S16**. Results epithelial transport assay

**Table S17**. Drug screen results gain of function library

**Table S18**. Characterized genes encoding for drug-metabolizing enzymes from bacteria

**Table S19**. Structurally refined strain activity prediction for BACUNI_RS05305

**Table S20**. Chemicals purchased for this study

**Table S21**. Plasmids used in this study

**Table S22**. Primers used in this study
